# Pharmacological METTL3 inhibition attenuates HIV-1 latency reversal in CD4^+^ T cells

**DOI:** 10.64898/2026.03.18.712554

**Authors:** Tarun Mishra, Avary Edwards, Li Wu

**Author notes:** Address correspondence to Li Wu.

## Abstract

*N*^6^-methyladenosine (m^6^A) is a major epitranscriptomic modification that regulates RNA metabolism and affects the replication and latency reversal of human immunodeficiency virus type 1 (HIV-1) in cells. Methyltransferase-like 3 (METTL3) is the principal catalytic enzyme responsible for m^6^A deposition, and its pharmacological inhibition has emerged as a potential therapeutic strategy for cancer and viral infections. However, the relative potency of METTL3 inhibitors in reducing m^6^A levels and their effects on HIV-1 latency reversal remain undefined. Here, we compared three commercially available METTL3 inhibitors (STM2457, STM3006, and STC-15) to evaluate their ability to reduce RNA m^6^A levels, suppress HIV-1 latency reversal, and affect cell viability in latently infected J-Lat cells and primary CD4^+^ T cells. In J-Lat cells, STM3006 and STC-15 were more potent than STM2457 in reducing RNA m^6^A levels at 24 and 48 hours post-treatment, as reflected by lower half-maximal inhibitory concentrations (IC_50_). However, STM3006 and STC-15 exhibited significant cytotoxicity at concentrations above 2 µM at 48 hours post-treatment, whereas STM2457 displayed minimal toxicity across all tested doses. In primary CD4^+^ T cells from three healthy donors, all three inhibitors reduced RNA m^6^A levels but induced greater cytotoxicity compared with J-Lat cells, with comparable effects at optimized concentrations. Notably, reduced RNA m^6^A levels correlated with diminished HIV-1 latency reversal in both J-Lat cells and a primary central memory CD4^+^ T cell model. Together, these findings demonstrate differential potency and cytotoxicity among METTL3 inhibitors and support a critical role for m^6^A RNA modification in regulating HIV-1 latency reversal.

## INTRODUCTION

Although combined antiretroviral therapy effectively suppresses HIV-1 replication to undetectable levels, the infection remains incurable due to the establishment of stable latent reservoirs (1). These reservoirs allow the virus to remain immunologically invisible, allowing it to persist despite lifelong antiretroviral therapy (2, 3). The maintenance of latency involves several mechanisms, including the sequestration of key host transcription factors and transcriptional repression of the proviral genome (4). Currently, two potential curative approaches are tested: the “block-and-lock,” which requires the enforcement of permanent epigenetic silencing of HIV-1 gene expression, and the “shock-and-kill,” which aims to reactivate latent proviruses so that infected cells can be recognized and eliminated by the host immune system (5, 6). A deeper understanding of the epitranscriptomic regulators that control latency reversal is essential for advancing these proposed cure strategies.

Eukaryotic RNA harbors more than 170 post-transcriptional modifications, with *N*^6^-methyladenosine (m^6^A) being the most abundant modification found on both coding and non-coding RNA (7–9). m^6^A modification is dynamically installed, removed, and interpreted by a set of cellular proteins termed writers, erasers, and readers, respectively. These proteins cooperatively regulate RNA metabolism, including splicing, stability, translation, and localization (10–12). The m^6^A write complex contains the catalytic subunit methyltransferase 3 (METTL3), whereas METTL14 primarily maintains the complex and facilitates binding of the substrate RNA (13, 14).

Many pathogenic viruses exploit the m^6^A RNA modification machinery to modulate host cellular processes, thereby optimizing viral replication and enhancing immune evasion and pathogenicity (15, 16). For example, HIV-1 exploits m^6^A modification to promote viral replication in cells and suppress innate immune responses (10, 17–22). Severe acute respiratory syndrome coronavirus 2 (SARS-CoV-2) relies on METTL3-driven m^6^A modification to stabilize viral RNAs and protect them from innate immune detection (23). Our published m^6^A profile of SARS-CoV-2-infected human lung epithelial cells also indicates that m^6^A modification of cellular RNA as a mechanism of posttranscriptional gene regulation during SARS-CoV-2 infection (24). SARS-CoV-2 and influenza A virus utilize METTL3-dependent m^6^A modification to support viral protein expression, regulate RNA splicing, and enhance viral replication (25, 26). Indeed, targeting the m^6^A modification pathway with a METTL3 inhibitor blocks replication of SARS-CoV-2 and a seasonal human β-coronavirus in cells (27), suggesting a novel approach to antiviral development. Given that HIV-1 infection and latency reversal enhance cellular RNA m^6^A levels (28–30), targeting METTL3 with inhibitors represents an effective therapeutic target. However, the relative potency of METTL3 inhibitors in reducing m^6^A levels and their effects on HIV-1 latency reversal remain undefined.

The methyltransferase reaction catalyzed by METTL3 depends on S-adenosyl-L-methionine (SAM) as a methyl donor, making the SAM-binding site an attractive target for small-molecule inhibitor design (31). These inhibitors can effectively reduce methyltransferase activity through competitive binding (9). The first-in-class METTL3 inhibitor STM2457 exhibits potent effects against myeloid leukemia in a mouse model (32). STC-15, a more potent derivative of STM2457, has been evaluated in clinical trials for advance cancer therapy, including a completed phase I (33, 34) and an ongoing phase Ib/II trial (https://clinicaltrials.gov/study/NCT06975293). Furthermore, STM3006, a second-generation METTL3 inhibitor with increased potency, has been developed as a potential cancer therapeutic (35).

In this study, we compared the efficacy of STM2457, STM3006, and STC-15 to evaluate their ability to reduce RNA m^6^A levels, suppress HIV-1 latency reversal, and affect cell viability in latently infected J-Lat cells and primary CD4^+^ T cells. Our findings indicate that RNA m^6^A modification plays a crucial role in regulating HIV-1 latency reversal in CD4^+^ T cells and underscore METTL3 inhibition as a potential strategy to modulate viral reactivation dynamics.

## RESULTS

### METTL3 inhibitors affect viability and reduce cellular RNA m^6^A levels in J-Lat 10.6 cells

To assess the effect of three METTL3 inhibitors, STM2457, STM3006, and STC-15 (summarized in Table 1) on CD4^+^ T cell viability and RNA m^6^A levels, HIV-1 latently infected J-Lat cells (clone 10.6) (36, 37) were treated with variable concentrations of each inhibitor for 24 h and 48 h. Treatment of cells with dimethyl sulfoxide (DMSO) was used as a vehicle control. First, cell viability was evaluated at 24 h and 48 h post-treatment. Compared with the DMSO-treated cells, STM2457 showed minimal cytotoxicity at 24 h and 48 h post-treatment, indicating that it was the least toxic compound tested. In contrast, STM3006 induced substantial cytotoxicity within 24 h at concentrations above 2 µM (Fig. 1A). While treatment with STC-15 did not affect cell viability significantly after 24 h, prolonged STC-15 exposure for 48 h resulted in significant cell death at concentrations above 4 µM (Fig. 1A-B). Thus, STM3006 and STC-15 were toxic to J-Lat 10.6 cells at concentrations above 2 μM, whereas STM2457 was relatively safe to cells.

**FIG 1.**
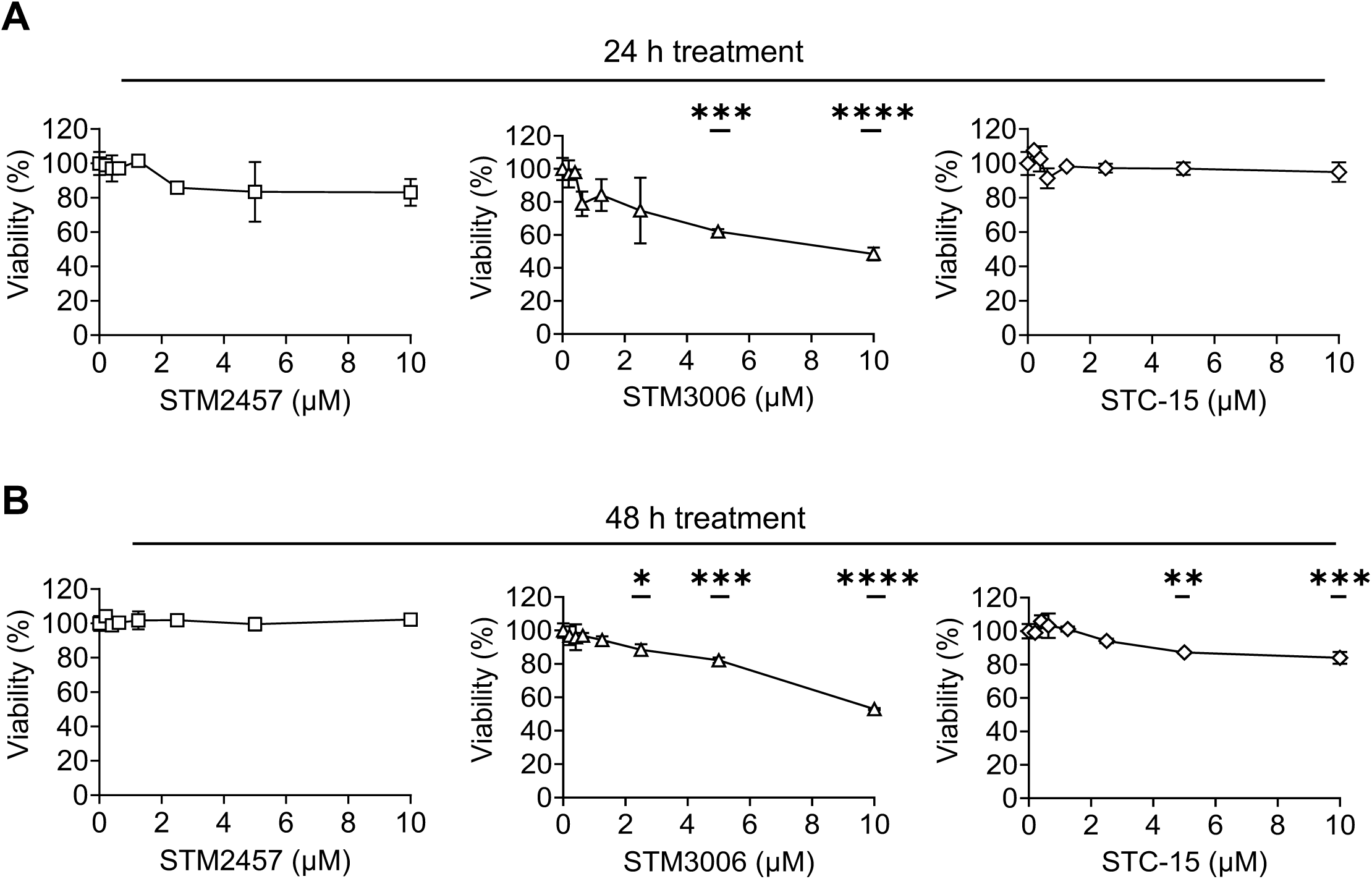
Evaluation of cell viability in METTL3 inhibitor-treated J-Lat 10.6 cells. The viability of J-Lat 10.6 cells was assessed by the MTT assay following treatment with DMSO (vehicle) or the indicated concentration of METTL3 inhibitors for (**A**) 24 h and (**B**) 48 h. Results are shown as mean ± SD from three independent experiments. Statistical analysis was performed using one-way ANOVA. * *P* < 0.05, ** *P* < 0.01, *** *P* < 0.001 and **** *P* < 0.0001. Statistical significance was compared between treated samples and the DMSO control.

**TABLE 1.**
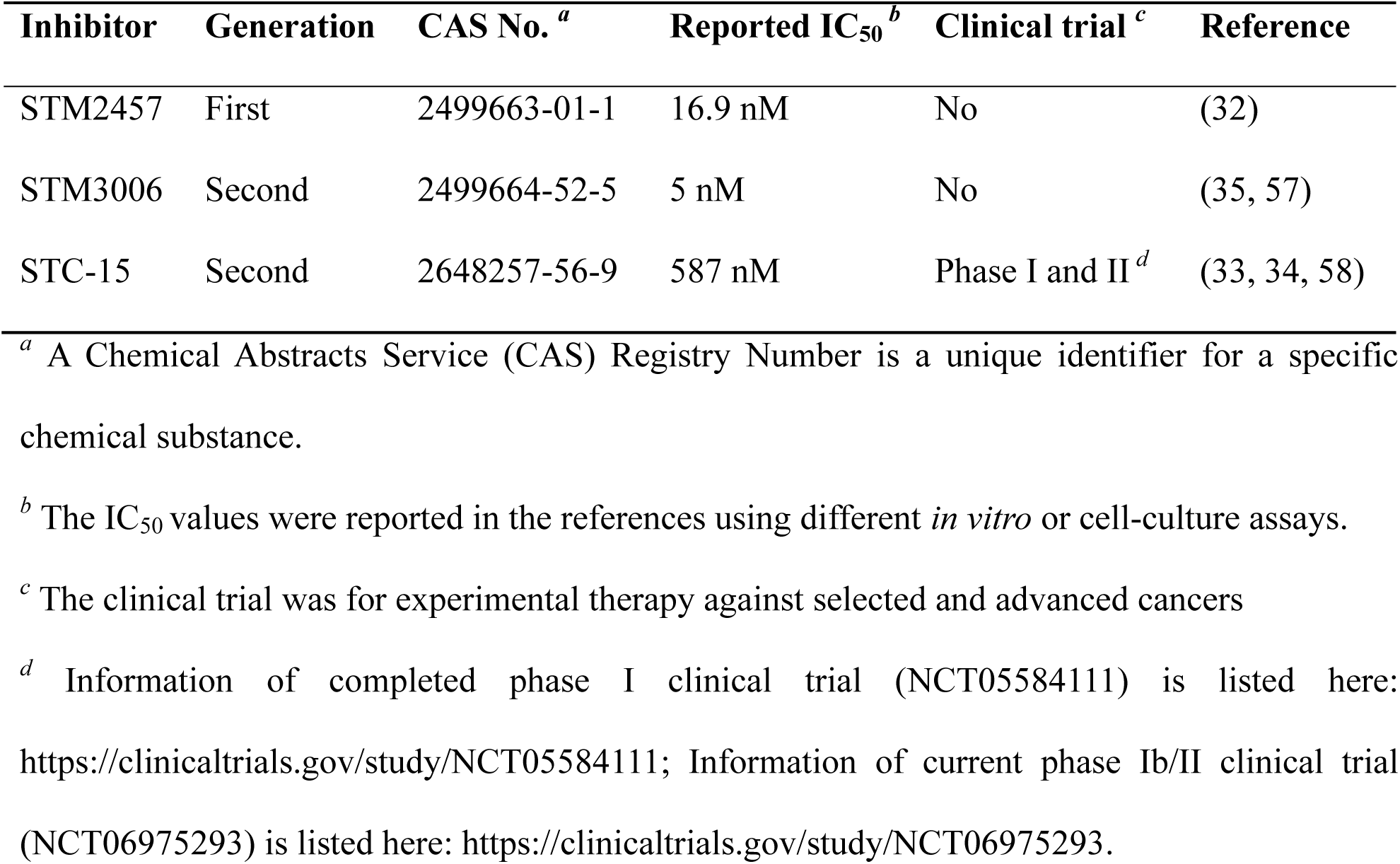
Summary of three METTL3-specific inhibitors.

| Inhibitor | Generation | CAS No. <sup>a</sup> | Reported IC<sub>50</sub> <sup>b</sup> | Clinical trial <sup>c</sup> | Reference |
| --- | --- | --- | --- | --- | --- |
| STM2457 | First | 2499663-01-1 | 16.9 nM | No | (32) |
| STM3006 | Second | 2499664-52-5 | 5 nM | No | (35, 57) |
| STC-15 | Second | 2648257-56-9 | 587 nM | Phase I and II <sup>d</sup> | (33, 34, 58) |

<sup>a</sup> A Chemical Abstracts Service (CAS) Registry Number is a unique identifier for a specific chemical substance.
<sup>b</sup> The IC<sub>50</sub> values were reported in the references using different *in vitro* or cell-culture assays.
<sup>c</sup> The clinical trial was for experimental therapy against selected and advanced cancers
<sup>d</sup> Information of completed phase I clinical trial (NCT05584111) is listed here: https://clinicaltrials.gov/study/NCT05584111; Information of current phase Ib/II clinical trial (NCT06975293) is listed here: https://clinicaltrials.gov/study/NCT06975293.

Next, to determine the optimal inhibitory concentration of the METTL3 inhibitors that reduce methyltransferase activity, total cellular RNA was isolated from DMSO- and METTL3 inhibitor-treated J-Lat 10.6 cells at 24 h and 48 h, and m^6^A levels were measured using enzyme-linked immunosorbent assay (ELISA) (36). At 24 h post-treatment, all three inhibitors reduced relative RNA m^6^A levels in a dose-dependent manner (Fig. 2). The mean ± SD of half-maximal inhibitory concentrations (IC_50_) were 3.86 ± 0.07 µM for STM2457, 1.44 ± 0.10 µM for STM3006, and 0.21 ± 0.09 µM for STC-15, demonstrating that STC-15 was the most potent inhibitor at the 24 h time point (Fig. 2A). At 48 h post-treatment, the IC_50_ values were 4.76 ± 0.10 µM for STM2457, 0.71 ± 0.08 µM for STM3006, and 0.75 ± 0.10 µM for STC-15. Thus, STM3006 and STC-15 exhibited comparable potency to reduce cellular RNA m^6^A levels, while STM2457 remained less effective (Fig. 2B). Together, these results indicate that STC-15 is the most effective inhibitor of METTL3 at 24 h post-treatment, whereas STM3006 and STC-15 showed similar efficacy after 48 h of treatment (Fig. 2A-B).

**FIG 2.**
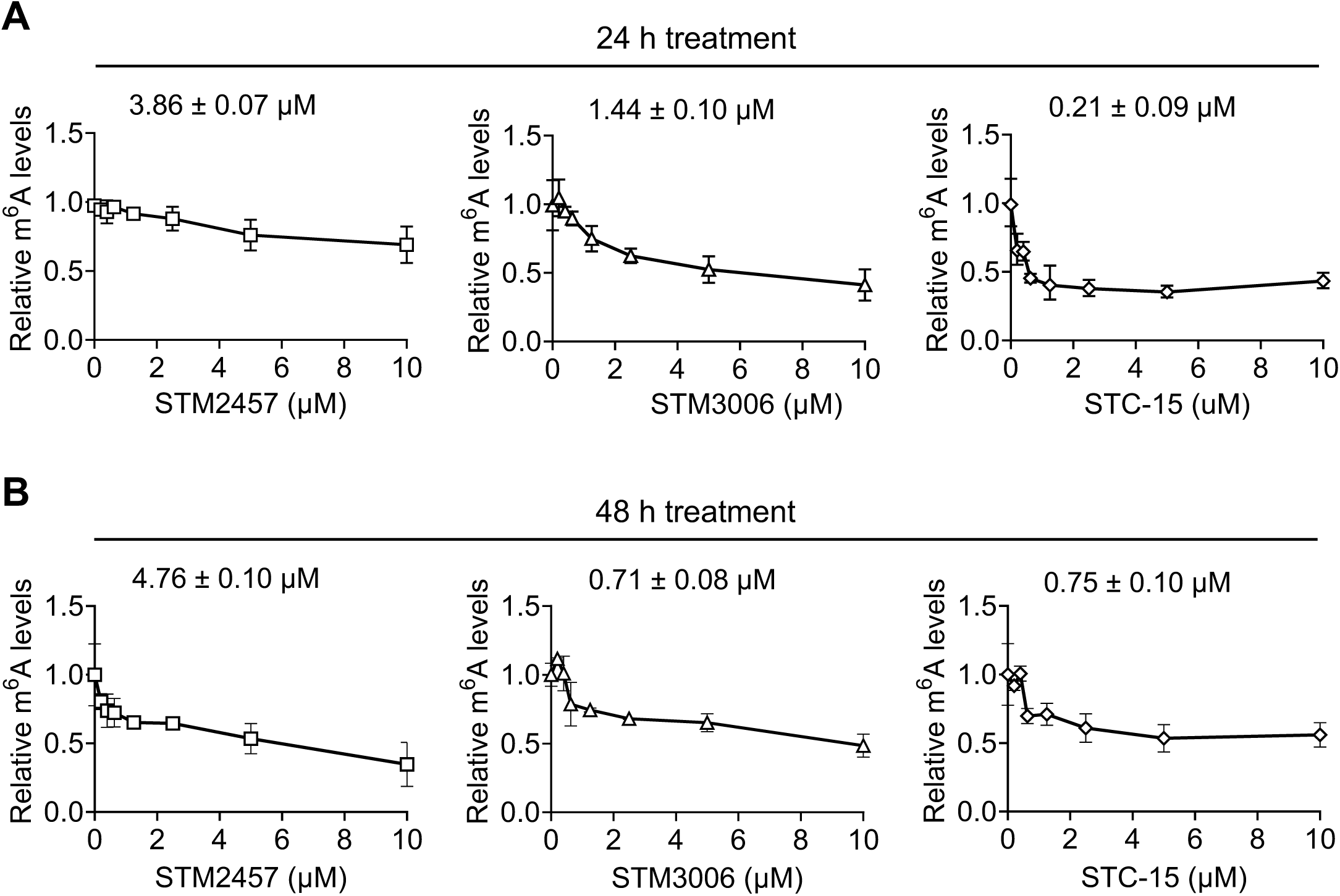
METTL3 inhibitors reduce the m^6^A levels of cellular RNA in J-Lat 10.6 cells. J-Lat 10.6 cells were treated with DMSO or the indicated concentration of METTL3 inhibitors. The m^6^A levels in total cellular RNA were quantified by ELISA following (**A**) 24 h or (**B**) 48 h treatment. The relative m^6^A levels were normalized to the DMSO control that was set as 1. The IC_50_ value of each inhibitor at 24 h and 48 h treatment is indicated on each figure panel.

### METTL3 inhibitors suppress HIV-1 latency reversal in J-Lat 10.6 cells

We have reported that inhibition of METTL3 activity using a distinct inhibitor UZH1a reduces HIV-1 latency reversal in latently infected J-Lat 10.6 cells (36). To extend these findings, we evaluated the effect of three more potent METTL3 inhibitors, STM2457, STM3006, and STC-15, on HIV-1 latency reversal in J-Lat 10.6 cells. Cells were treated with DMSO or each inhibitor with variable concentrations for 24 h or 48 h, followed by treatment with phorbol 12-myristate 13-acetate (PMA) and ionomycin (P+I) for 6 h to induce HIV-1 latency reversal, which was quantified by measuring green fluorescent protein (GFP) reporter expression using flow cytometry (36). All three METTL3 inhibitors significantly reduced P+I-induced GFP expression compared to DMSO-treated control at both 24 h and 48 h (Fig. 3A-B). Among the inhibitors tested, STM3006 consistently produced the greatest reduction in GFP expression, indicating the strongest suppression of latency reversal. STM2457 exhibited intermediate potency, whereas STC-15 showed the least inhibitory effect (Fig. 3A-B). Collectively, these results demonstrate that pharmacological inhibition of METTL3 suppresses HIV-1 latency reversal in J-Lat 10.6 cells, with relative potency ranked as follows: STM3006 > STM2457 > STC-15 (Fig. 3A-B).

**FIG 3.**
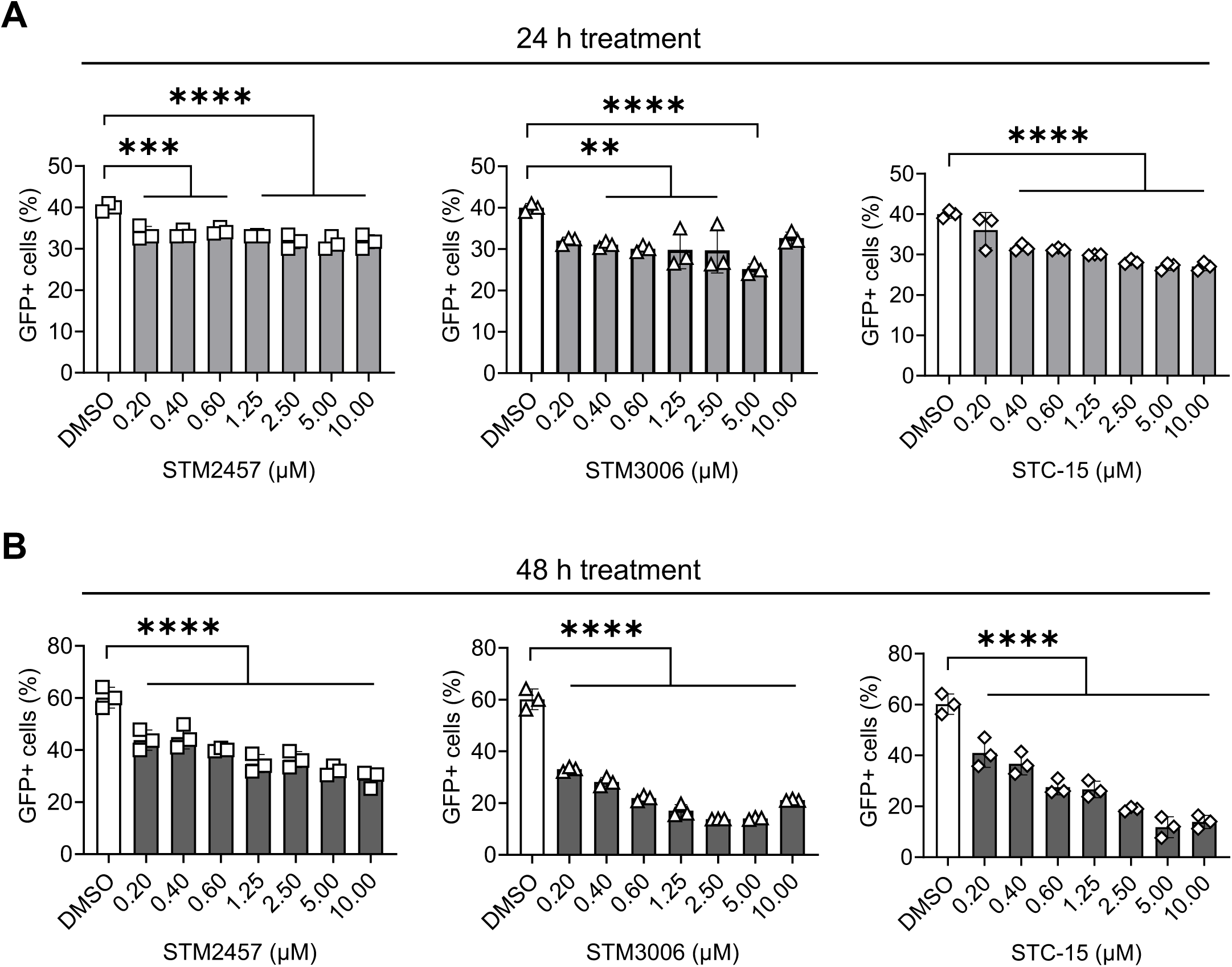
Inhibition of METTL3 activity reduces HIV-1 latency reversal. J-Lat 10.6 cells were treated with DMSO or the indicated concentration of METTL3 inhibitors for (**A**) 24 h or (**B**) 48 h, followed by stimulation with P+I for 6 h. The percentage of GFP-positive cells was determined by flow cytometry after the 6 h P+I treatment. Data are presented as mean ± SD from three independent experiments. Statistical analysis was performed using one-way ANOVA. ** *P* < 0.01, *** *P* < 0.001, and **** *P* < 0.0001. Statistical significance was compared between treated samples and the DMSO control.

### METTL3 inhibitors affect viability and reduce RNA m^6^A levels in primary CD4^+^ T cells

To determine whether the effects observed in J-Lat 10.6 cells extend to physiologically relevant primary T cells, we next evaluated the impact of METTL3 inhibition on the viability of primary CD4^+^ T cells and RNA m^6^A levels. Primary CD4^+^ T cells from three independent healthy donors were treated with DMSO or with optimized concentrations of STM2457, STM3006, and STC-15 for 24 h and 48 h (Fig. 4A-B). The inhibitor concentrations were selected based on the viability profile observed in J-Lat 10.6 cells (Fig. 1A-B). Cell viability was measured at 24 h and 48 h post-treatment. Primary CD4^+^ T cells exhibited significant reductions in viability at both time points across all tested concentrations compared to DMSO-treated controls (Fig. 4A-B), suggesting that primary CD4^+^ T cells are more sensitive to METTL3 inhibitor treatment than J-Lat 10.6 cells.

**FIG 4.**
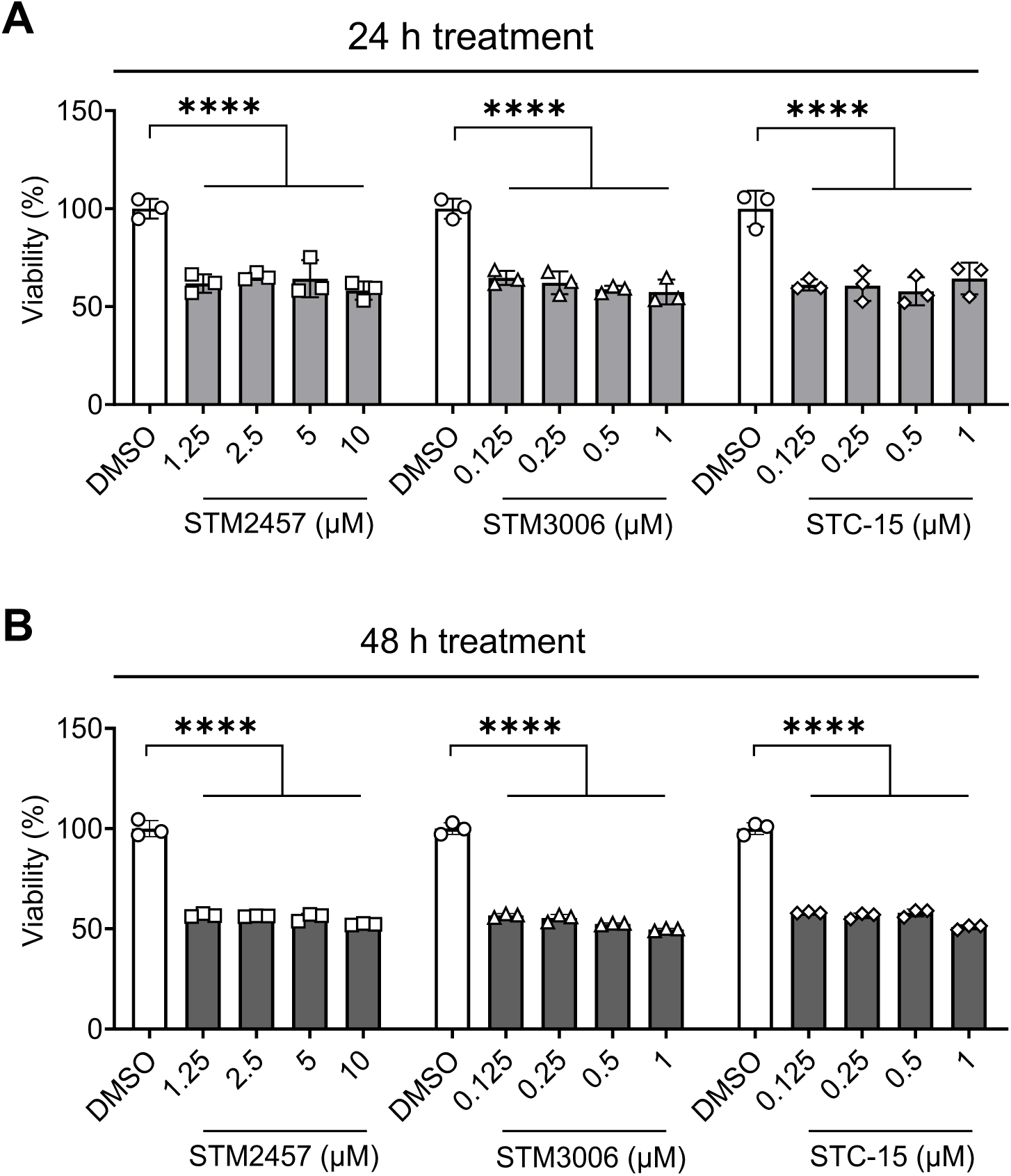
Evaluation of cell viability in METTL3 inhibitor-treated primary CD4^+^ T cells. The viability of primary CD4^+^ T cells was assessed by the MTT assay following treatment with DMSO or the indicated concentration of METTL3 inhibitors for (**A**) 24 h and (**B**) 48 h. Results are shown as mean ± SD from three independent donor experiments. Statistical analysis was performed using two-way ANOVA. * *P* < 0.05, ** *P* < 0.01, *** *P* < 0.001 and **** *P* < 0.0001. Statistical significance was compared between treated samples and the DMSO control.

We next assessed the dose effect of METTL3 inhibitors on RNA m^6^A levels in primary CD4^+^ T cells from three independent healthy donors. Total cellular RNA m^6^A was isolated after 24 h and 48 h of treatment, and RNA m^6^A levels were quantified. STM2457 and STM3006 induced a consistent dose-dependent reduction in RNA m^6^A Levels in cells from all three donors at both time points (Fig. 5A-B). In contrast, STC-15 demonstrated a comparatively weaker effect in reducing m^6^A levels in primary CD4^+^ T cells under the tested conditions (Fig. 5A-B). At 24 h post-treatment, the IC_50_ values (mean ± SD) were 0.93 ± 0.25 µM for STM2457, 0.61 ± 0.21 µM for STM3006, and 2.47 ± 0.23 µM for STC-15. At 48 h post-treatment, the IC_50_ values were 2.98 ± 0.24 µM for STM2457, 0.81 ± 0.17 µM for STM3006, and 4.04 ± 0.21 µM for STC-15. Thus, STM3006 was the most potent inhibitor to reduce RNA m^6^A levels in primary CD4^+^ T cells at both 24 and 48 h post-treatment. These findings indicate that STM2457 and STM3006 are more effective than STC-15 in reducing RNA m^6^A levels in primary CD4^+^ T cells, highlighting cell-type-specific differences in response to METTL3 inhibition compared with J-Lat 10.6 cells (Fig. 5 vs. Fig. 2).

**FIG 5.**
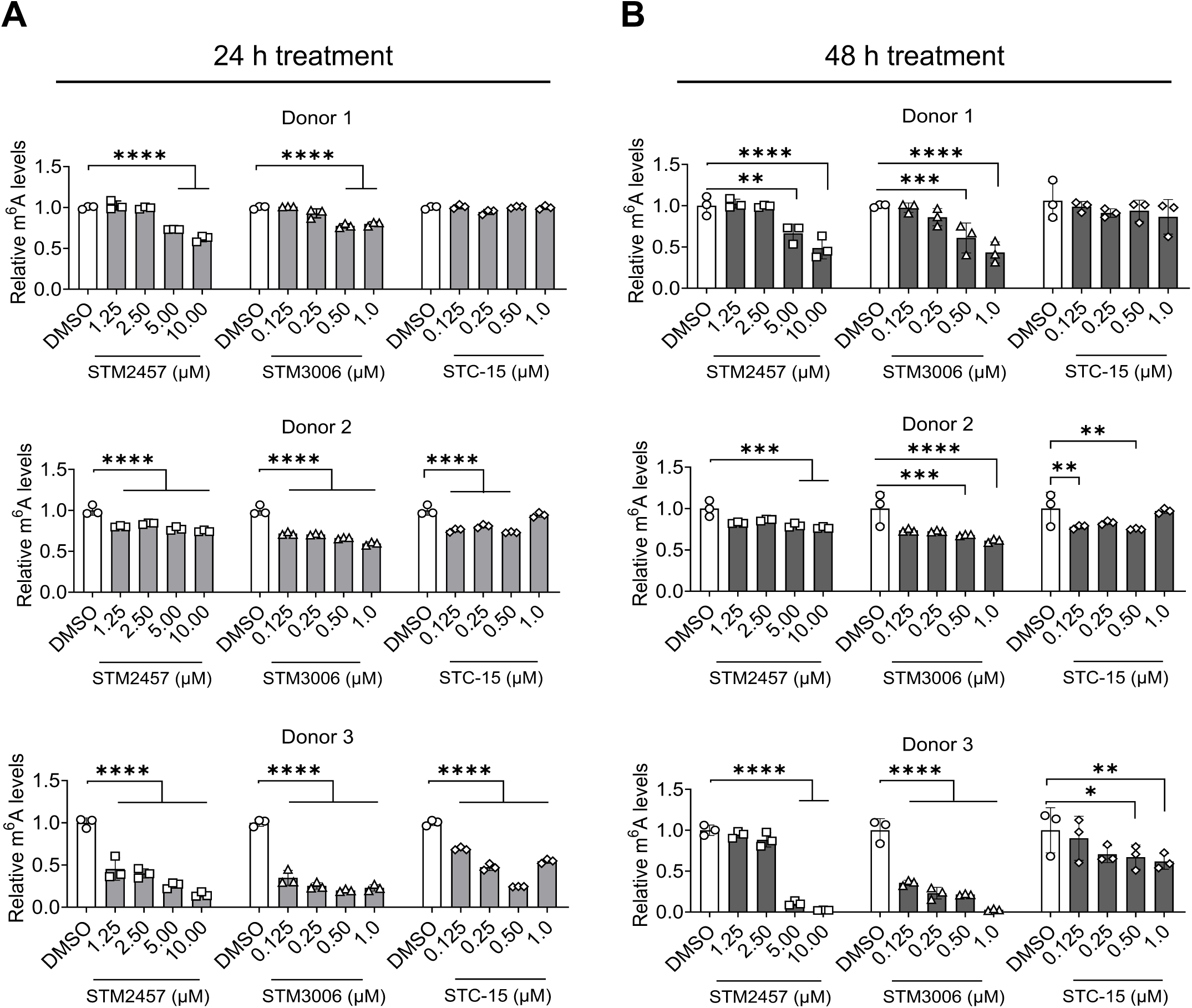
METTL3 inhibitors reduce m^6^A levels of cellular RNA in primary CD4⁺ T cells. Primary CD4⁺ T cells isolated from three independent donors were treated with DMSO or the indicated concentrations of METTL3 inhibitors. The m^6^A levels in total cellular RNA were quantified by ELISA after (**A**) 24 h or (**B**) 48 h of treatment. The relative m^6^A levels were normalized to the DMSO control that was set as 1. Bar graphs represent data from three independent donors treated with the indicated concentrations of METTL3 inhibitors. Statistical analysis was performed using two-way ANOVA. * *P* < 0.05, ** *P* < 0.01, *** *P* < 0.001, and **** *P* < 0.0001. Statistical significance was compared between treated samples and the DMSO control.

### METTL3 inhibitors reduce HIV-1 latency reversal in primary central memory CD4^+^ T cells (T_CM_)

To examine the effects of METTL3 inhibitors on HIV-1 latency reversal, we employed an *ex vivo* primary T_CM_ model of HIV-1 latency using our optimized protocol (36, 38) as outlined in Fig. 6A. HIV-1 GFP expression of T_CM_ was assessed on day 10 and day 22 after transduction with a GFP reporter HIV-1 vector on day 7 to confirm productive infection and the establishment of latency, respectively (Fig. 6A-B). On day 10, approximately 12%-15% of CD4^+^ T cells from three donors expressed GFP, indicating productive infection (Fig. 6B and data not shown). On day 17, cells were treated with either DMSO (vehicle control) or STM2457 (10 µM), STM3006 (1 µM), or STC-15 (1 µM) for two days. Two days post-treatment, cells were reactivated using a P+I combination in the presence of separate METTL3 inhibitors. GFP expression was quantified as a measure of HIV-1 latency reversal three days post-reactivation on day 22. Medium-treated cells served as a negative control and showed basal GFP expression levels ranging from 0.6% to 1% (Fig. 6B and data not shown). Compared to the DMSO control, cellular RNA m^6^A levels were significantly reduced with treatment of all three METTL3 inhibitors by 2-fold on day 22 (Fig. 6C). Furthermore, treatment with all three METTL3 inhibitors reduced HIV-1 reactivation by approximately 2.2-fold compared with DMSO-treated T_CM_ across three donors, as measured by both the percentage and mean fluorescence intensity (MFI) of GFP-positive cells (Fig. 6D-E). These findings indicate that reducing RNA m^6^A levels with METTL3 inhibitor treatment attenuates HIV-1 reactivation in latently infected primary CD4^+^ T cells.

**FIG 6.**
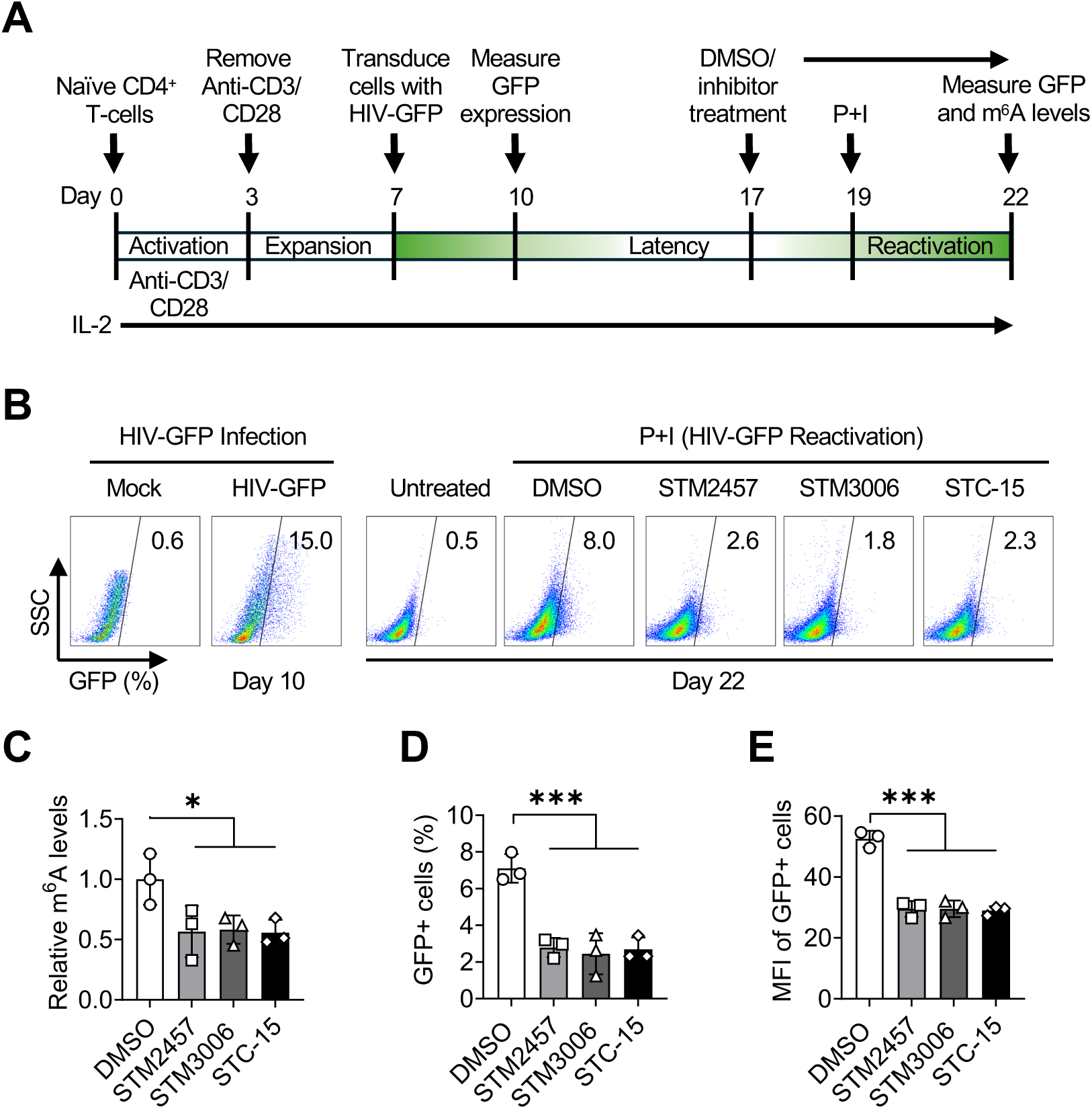
Treatment with METTL3 inhibitors reduces HIV-1 latency reversal in primary CD4^+^ T_CM_ cells. (**A**) Protocol summary. Naïve CD4^+^ T cells were isolated from PBMCs of three healthy donors. Cells were activated using anti-CD3/CD28 antibody-coated beads for 3 days and infected with single-cycle HIV-1-GFP reporter at day 7 to generate a primary T_CM_ model of latency. (**B**) GFP expression was measured at day 10 and 22 by flow cytometry. Representative data from a single donor are shown. SSC, side scatter. The percentage of GFP-positive cells are shown on the plots. (**C**) After 10 days of HIV-1 infection and culture, cells were treated with either DMSO or METTL3 inhibitors and m^6^A levels of cellular RNA were measured by ELISA. The relative m^6^A levels were normalized to the DMSO control that was set as 1. Results from three individual donors are shown. (**D** and **E**) HIV-1 latency reversal was measured by GFP expression using flow cytometry at day 22. Changes in (D) GFP percentage and (E) mean fluorescence intensity (MFI) of GFP-positive cells from three independent donors are shown. Statistical analysis was performed using two-way ANOVA. * *P* < 0.05, and *** *P* < 0.001.

## DISCUSSION

Despite the progress of antiretroviral therapy, the most challenging obstacle in HIV-1 cure is the persistence of latently infected resting CD4^+^ T cells (39, 40). These cells carry transcriptionally inactive proviruses that can fully reactivate and produce an infectious virus (41). Therefore, a better understanding of the molecular mechanism that controls latency reactivation is essential for achieving a functional cure. In this study, we investigated the role of m^6^A RNA modification in HIV-1 latency reversal using three commercially available METTL3 inhibitors for their ability to attenuate HIV-1 reactivation in latently infected J-Lat 10.6 cells and primary central memory CD4^+^ T cells. Consistent with previous studies using knockdown of METTL3 or activators of the METTL3 complex to modulate HIV-1 replication and gene expression (42, 43), our results demonstrate that inhibition of METTL3 suppresses HIV-1 latency reversal in both J-Lat 10.6 cells and a primary T_CM_ latency model. These data provide functional evidence linking m^6^A modification to the control of latent viral RNA expression in both transformed CD4^+^ T cell lines and primary CD4^+^ T cells.

Considering the importance of m^6^A RNA modification in normal cell growth and physiology, the potential side effects of METTL3 inhibitors on cell viability must be monitored (44). Notably, we observed variable cytotoxicity among the three inhibitors. While STM2457 showed minimal cytotoxicity, STM3006 and STC-15 reduced cell viability in J-Lat 10.6 cells, consistent with prior studies demonstrating that STM2457 is relatively safe at concentrations below 32 μM (45). While STM3006 and STC-15 are known to exhibit antiproliferative effects, they induce apoptosis and suppress growth in cancer cells by interferon induction through double-stranded RNA formation (35, 46, 47). The elevated sensitivity of primary CD4⁺ T cells to METTL3 inhibition suggests that methyltransferase activity is critical for T cell survival and RNA homeostasis, as METTL3 has been implicated in maintaining mRNA stability, splicing, and translational efficiency in numerous cell types (9, 48–50).

Among the inhibitors, STM3006 displayed the strongest ability to suppress HIV-1 latency reversal and reduce cellular RNA m^6^A levels, followed by STM2457 and STC-15. These results confirm that the inhibitors functionally target the m^6^A modification pathways and demonstrate that METTL3 enzymatic activity contributes to maintaining m^6^A levels in CD4⁺ T cells. This observation also validates that downstream effects on HIV-1 latency reversal are linked to altered epitranscriptomic regulation (10, 36).

Mechanistically, the reduction in HIV-1 reactivation may arise from multiple layers of post-transcriptional control. It has been shown that m^6^A modification regulates viral RNA export from the nucleus by recruiting m^6^A reader proteins that facilitate nuclear-cytoplasmic trafficking and the stability of transcripts of HIV-1 and hepatitis B virus (42, 51). Similar m^6^A-dependent mechanisms have been reported in other viruses, such as hepatitis B virus and the Zika virus, in which methylation enhances viral mRNA translation and replication (52, 53). Therefore, suppression of this pathway via METTL3 inhibition could broadly reduce viral RNA stability, splicing efficiency, or translational output, collectively diminishing productive viral infection or reactivation.

From a therapeutic standpoint, these results have implications for HIV-1 latency management strategies. While “shock-and-kill” approaches aim to reactivate and eliminate latent HIV-1 reservoirs (54), our findings point toward an alternative “block-and-lock” model, in which inhibition of m^6^A modification restricts latency reactivation and maintains proviruses in latent, non-replicative state (36, 43, 55). However, the observed cytotoxicity of METTL3 inhibitors, particularly in primary CD4⁺ T cells, demands the need for more selective, less toxic analogs or delivery strategies that minimize off-target effects. Given the central role of METTL3 in cellular RNA metabolism, any therapeutic application must carefully balance antiviral efficacy with host cell viability.

In summary, our study establishes that inhibition of METTL3-mediated m^6^A methylation suppresses HIV-1 latency reversal in both J-Lat 10.6 and T_CM_ models, thereby linking epitranscriptomic regulation to HIV-1 reactivation. Among the tested three METTL3-specific inhibitors, STM3006 displayed the highest potency but also significant cytotoxicity. These results highlight the dual nature of METTL3 inhibition: while it can effectively block viral reactivation, it also disrupts essential host RNA processing functions. Further research using latently infected cells from HIV-1 patients and next-generation METTL3 modulators will be crucial to delineate the therapeutic potential of targeting RNA methylation in HIV-1 latency.

## MATERIALS AND METHODS

### Cell culture

J-Lat 10.6 cells (37) were obtained from the NIH AIDS Reagent Program, Division of AIDS, NIAID, NIH. Primary CD4^+^ T cells were isolated from healthy donor blood obtained from the DeGowin Blood Center at the University of Iowa using the EasySep Human CD4^+^ T cell isolation kit (STEMCELL Technologies). J-Lat 10.6 and primary CD4^+^ T cells were cultured in RPMI-1640 (ATCC) supplemented with 10% fetal bovine serum (FBS; R&D systems) and antibiotics (100 U/mL penicillin and 100 µg/mL streptomycin, Gibco). All cells were cultured in 5% CO_2_ at 37°C (3, 36). Cells used in the study tested negative for mycoplasma contamination using a PCR-based universal mycoplasma detection kit (ATCC 30-1012K).

### METTL3 inhibitors treatment and cell viability assay

The J-Lat 10.6 cells and primary CD4^+^ T cells were seeded at a concentration of 5 × 10^5^ /mL 24 h before treatment with DMSO (Sigma-Aldrich) or METTL3 inhibitors STM2457 (Cat. No. HY-134836), STM3006 (Cat. No. HY-156773), and STC-15 (Cat. No. HY-156677) that were purchased from MedChemExpress. The cells were treated with inhibitor concentrations ranging from 0.20 µM to 10 µM, as indicated in the figures and legends. Viability was assessed by using the CellTitre 96 Non-radioactive Cell proliferation assay (MTT assay, Promega) as described (36).

### m^6^A quantification by ELISA

Total cellular RNA was isolated using the RNeasy Plus Kit (Qiagen) following the manufacturer’s protocol. The RNA was quantified using the Nanodrop-OneC spectrophotometer (Thermo Fisher). The RNA m^6^A modification levels were quantified with 200 ng of total cellular RNA per sample using m^6^A ELISA as described (28, 36, 56).

### HIV- 1 latency reversal and flow cytometry

J-Lat 10.6 cells (5 × 10^6^) were seeded in T75 flasks 24 h before treatment with DMSO or various concentrations of STM2457, STM3006, and STC-15. After 24 h and 48 h of inhibitor treatment, cells were stimulated with 32 nM PMA (Sigma, P1585) and 1 µM ionomycin (Sigma, I0634) for 6 h. Cells were then fixed with 4% paraformaldehyde (36). The percentage of GFP-positive cells was determined by flow cytometry and analyzed by FlowJo software.

### T_CM_ model of HIV-1 latency and reactivation

The T_CM_ latency model was performed as previously described (36, 38). Briefly, naïve CD4^+^ T cells were isolated from healthy donor blood using the EasySep Human CD4^+^ T cell isolation kit (STEMCELL Technologies). Naïve CD4^+^ T cells were activated with CD3/CD28 Dynabeads for 3 days, rested for 4 days, then infected with (1 × 10^6^ cells) stomatitis virus G protein (VSV-G)-pseudotyped single-cycle HIV-1 reporter (NL4-3ΔEnv-GFP). Cells were cultured for an additional 7 days to establish latency. Latently infected cells were treated either with DMSO, STM2457 (10 µM), STM3006 (1 µM), or STC-15 (1 µM) for the next 2 days, followed by P+I combination-mediated activation for 3 days. The METTL3 inhibitors were replenished on day 19. On day 22, total RNA m^6^A levels were measured by ELISA, and GFP expression representing HIV-1 latency reversal was assessed by flow cytometry.

## ACKNOWLEDGMENTS

This work was supported by the U.S. National Institutes of Health, grant number R33AI169659 to L.W. L.W. is also supported by NIH grants R01AI189220, R21AI181742, and P30CA086862 supplement. The content is solely the responsibility of the authors and does not necessarily represent the official views of the NIH. J-Lat 10.6 cell clone was obtained from NIH AIDS reagent program, Division of AIDS, NIAID, and NIH. We thank the members of the Wu lab for helpful discussions and suggestions.

## DATA AVAILABILITY

All data and materials are included in the manuscript.

## AUTHOR CONTRIBUTIONS

Conceptualization: T.M., and L.W. Resources: L.W. Methodology, investigation, and validation: T.M., and A.E. Formal analysis: T.M., A.E., and L.W. Writing-original draft: T.M. and L.W. Writing-review and editing: T.M., and L.W. Visualization: T.M. and L.W. Supervision, Project administration, and Funding acquisition: L.W.

